# Pcadapt: An R Package to Perform Genome Scans for Selection Based on Principal Component Analysis

**DOI:** 10.1101/056135

**Authors:** Keurcien Luu, Eric Bazin, Michael G. B. Blum

**Affiliations:** Université Grenoble Alpes, CNRS, Laboratoire TIMC-IMAG, UMR 5525, France; Université Grenoble Alpes, CNRS, Laboratoire d’Ecologie Alpine UMR 5553, France

**Keywords:** population genetics, R package, outlier detection, Mahalanobis distance, principal component analysis

## Abstract

The R package *pcadapt* performs genome scans to detect genes under selection based on population genomic data. It assumes that candidate markers are outliers with respect to how they are related to population structure. Because population structure is ascertained with principal component analysis, the package is fast and works with large-scale data. It can handle missing data and pooled sequencing data. By contrast to population-based approaches, the package handle admixed individuals and does not require grouping individuals into populations. Since its first release, *pcadapt* has evolved both in terms of statistical approach and software implementation. We present results obtained with robust Mahalanobis distance, which is a new statistic for genome scans available in the 2.0 and later versions of the package. When hierarchical population structure occurs, Mahalanobis distance is more powerful than the communality statistic that was implemented in the first version of the package. Using simulated data, we compare *pcadapt* to other software for genome scans (*BayeScan, hapflk, OutFLANK, sNMF*). We find that the proportion of false discoveries is around a nominal false discovery rate set at 10% with the exception of *BayeScan* that generates 40% of false discoveries. We also find that the power of *BayeScan* is severely impacted by the presence of admixed individuals whereas *pcadapt* is not impacted. Last, we find that *pcadapt* and *hapflk* are the most powerful software in scenarios of population divergence and range expansion. Because *pcadapt* handles next-generation sequencing data, it is a valuable tool for data analysis in molecular ecology.

## Introduction

Looking for variants with unexpectedly large differences of allele frequencies between populations is a common approach to detect signals of natural selection (Lewontin and Krakauer, 1973). When variants confer a selective advantage in the local environment, allele frequency changes are triggered by natural selection leading to unexpectedly large differences of allele frequencies between populations. To detect variants with large differences of allele frequencies, numerous test statistics have been proposed, which are usually based on chi-square approximations of *F_ST_*-related test statistics (François et al., 2016).

Statistical approaches for detecting selection should address several challenges. The first challenge is to account for hierarchical population structure that arises when genetic differentiation between populations is not identical between all pairs of populations. Statistical tests based on *F_ST_* that do not account for hierarchical structure, when it occurs, generate a large excess of false positive loci (Bierne et al., 2013; Excoffier et al., 2009).

A second challenge arises because approaches based on *F_ST_*-related measures require to group individuals into populations, although defining populations is a difficult task (Waples and Gaggiotti, 2006). Individual sampling may not be population-based but based on more continuous sampling schemes (Lotterhos and Whitlock, 2015). Additionally assigning an admixed individual to a single population involves some arbitrariness because different regions of its genome might come from different populations (Pritchard et al., 2000). Several individual-based methods of genome scans have already been proposed to address this challenge and they are based on related techniques of multivariate analysis including principal component analysis (PCA), factor models, and non-negative matrix factorization (Duforet-Frebourg et al., 2014; Hao et al., 2016; Galinsky et al., 2016; Chen et al., 2016; Duforet-Frebourg et al., 2016; Martins et al., 2016).

The last challenge arises from the nature of multilocus datasets generated from next generation sequencing platforms. Because datasets are massive with a large number of molecular markers, Monte Carlo methods usually implemented in Bayesian statistics may be prohibitively slow (Lange et al., 2014). Additionally, next generation sequencing data may contain a substantial proportion of missing data that should be accounted for (Arnold et al., 2013; Gautier et al., 2013).

To address the aforementioned challenges, we have developed the software *pcadapt* and the R package *pcadapt*. The software *pcadapt* is now deprecated and the R package only is maintained. *pcadapt* assumes that markers excessively related with population structure are candidates for local adaptation. Since its first release, *pcadapt* has substantially evolved both in terms of statistical approach and software implementation (Table 1).

**Table 1.** Summary of the different statistical methods and implementations of *pcadapt*. Pop. structure stands for population structure and dist. stands for distance.

| Test statistic | Pop. structure | Language | Command line | Versions of the R package | Ref. |
| --- | --- | --- | --- | --- | --- |
| Bayes factor | Factor model | C | PCAdapt | NA | Duforet-Frebourg et al. (2014) |
| Communality | PCA | C and R | PCAdapt fast | 1.x | Duforet-Frebourg et al. (2016) |
| Mahalanobis dist. | PCA | R | NA | 2.x and 3.x | This paper |

The first release of *pcadapt* was a command line C software. It implemented a Monte Carlo approach based on a Bayesian factor model (Duforet-Frebourg et al., 2014). The test statistic for outlier detection was a Bayes factor. Because Monte Carlo methods can be computationally prohibitive with massive NGS data, we then developed an alternative approach based on PCA. The first statistic based on PCA was the *communality* statistic, which measures the percentage of variation of a SNP explained by the first *K* principal components (Duforet-Frebourg et al., 2016). It was initially implemented with a command-line C software (the *pcadapt fast* command) before being implemented in the *pcadapt* R package. We do not maintain C versions of *pcadapt* anymore. The whole analysis that goes from reading genotype files to detecting outlier SNPs can now be performed in R (R Core Team, 2015).

The 2.0 and following versions of the R package implement a more powerful statistic for genome scans. The test statistic is a robust Mahalanobis distance. A vector containing *K z*-scores measures to what extent a SNP is related to the first *K* principal components. The Mahalanobis distance is then computed for each SNP to detect outliers for which the vector of *z*-scores do not follow the distribution of the main bulk of points. The term robust refers to the fact that the estimators of the mean and of the covariance matrix of *z*, which are required to compute Mahalanobis distances, are not sensitive to the presence of outliers in the dataset (Maronna and Zamar, 2012). In the following, we provide a comparison of statistical power that shows that Mahalanobis distance provides more powerful genome scans compared to the communality statistic and to the Bayes factor that were implemented in previous versions of *pcadapt*.

In addition to comparing the different test statistics that were implemented in *pcadapt*, we compare statistic performance obtained with the 3.0 version of *pcadapt* and with other software of genome scans. We use simulated data to compare software in terms of false discovery rate (FDR) and statistical power. We consider data simulated under different demographic models including island model, divergence model and range expansion. To perform comparisons, we include software that require to group individuals into populations: *BayeScan* (Foll and Gaggiotti, 2008), the *F_LK_* statistic as implemented in the *hapflk* software (Bonhomme et al., 2010), and *OutFLANK* that provides a robust estimation of the null distribution of a *F_ST_* test statistic (Whitlock and Lotterhos, 2015). We additionally consider the *sNMF* software that implements another individual-based test statistic for genome scans (Frichot et al., 2014; Martins et al., 2016).

## Statistical and Computational approach

### Input data

The R package can handle different data formats for the genotype data matrix. In the version 3.0 that is currently available on CRAN, the package can handle genotype data files in the *vcf*, *ped* and *lfmm* formats. In addition, the package can also handle a *pcadapt* format, which is a text file where each line contains the allele counts of all individuals at a given locus. When reading a genotype data matrix with the *read.pcadapt* function, a *.pcadapt* file is generated, which contains the genotype data in the *pcadapt* format.

### Choosing the number of principal components

In the following, we denote by *n* the number of individuals, by *p* the number of genetic markers, and by *G* the genotype matrix that is composed of *n* lines and *p* columns. The genotypic information at locus *j* for individual i is encoded by the allele count *G_ij_*, 1 ≤ *i* ≤ *n* and 1 ≤ *j* ≤ *p*, which is a value in 0, 1 for haploid species and in 0, 1, 2 for diploid species.

First, we normalize the genotype matrix columnwise. For diploid data, we consider the usual normalization in population genomics where 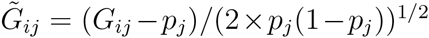, and *P_j_* denotes the minor allele frequency for locus *j* (Patterson et al., 2006). The normalization for haploid data is similar except that the denominator is given by (*p_j_*(1 − *p_j_*))^1/2^.

Then, we use the normalized genotype matrix 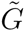 to ascertain population structure with PCA (Patterson et al., 2006). The number of principal components to consider is denoted *K* and is a parameter that should be chosen by the user. In order to choose *K*, we recommend to consider the graphical approach based on the scree plot (Jackson, 1993). The scree plot displays the eigenvalues of the covariance matrix Ω in descending order. Up to a constant, eigenvalues are proportional to the proportion of variance explained by each principal component. The eigenvalues that correspond to random variation lie on a straight line whereas the ones corresponding to population structure depart from the line. We recommend to use Cattell’s rule that states that components corresponding to eigenvalues to the left of the straight line should be kept (Cattell, 1966).

### Test statistic

We now detail how the package computes the test statistic. We consider multiple linear regressions by regressing each of the *p* SNPs by the *K* principal components *X*_1_, …, *X_K_*

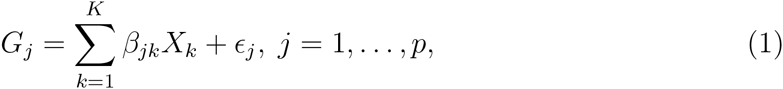

where *β_jk_* is the regression coefficient corresponding to the *j*-th SNP regressed by the *k*-th principal component, and *ϵ_j_* is the residuals vector. To summarize the result of the regression analysis for the *j*-th SNP, we return a vector of *z*-scores *z_j_* = (*z*_*j*1_, …, *z_jK_*) where *z_jk_* corresponds to the z-score obtained when regressing the *j*-th SNP by the *k*-th principal component.

The next step is to look for outliers based on the vector of *z*-scores. We consider a classical approach in multivariate analysis for outlier detection. The test statistic is a robust Mahalanobis distance *D* defined as

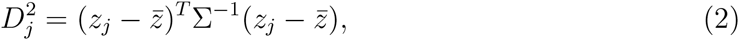

where Σ is the (*K* × *K*) covariance matrix of the *z*-scores and 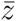 is the vector of the *K z*-score means (Maronna and Zamar, 2012). When *K* > 1, the covariance matrix Σ is estimated with the Orthogonalized Gnanadesikan-Kettenring method that is a robust estimate of the covariance able to handle large-scale data (Maronna and Zamar, 2012) *(covRob* function of the *robust* R package). When *K* =1, the variance is estimated with another robust estimate (*cov.rob* function of the *MASS* R package).

### Genomic Inflation Factor

To perform multiple hypothesis testing, Mahalanobis distances should be transformed into p-values. If the *z*-scores were truly multivariate Gaussian, the Mahalanobis distances D should be chi-square distributed with *K* degrees of freedom. However, as usual for genome scans, there are confounding factors that inflate values of the test statistic and that would lead to an excess of false positives (François et al., 2016). To account for the inflation of test statistics, we divide Mahalanobis distances by a constant λ to obtain a statistic that can be approximated by a chi-square distribution with *K* degrees of freedom. This constant is estimated by the genomic inflation factor defined here as the median of the Mahalanobis distances divided by the median of the chi-square distribution with *K* degrees of freedom (Devlin and Roeder, 1999).

### Control of the false discovery rate (FDR)

Once *p*-values are computed, there is a problem of decision-making related to the choice of a threshold for *p*-values. We recommend to use the FDR approach where the objective is to provide a list of candidate genes with an expected proportion of false discoveries smaller than a specified value. For controlling the FDR, we consider the *q*-value procedure as implemented in the *qvalue* R package that is less conservative than Bonferroni or Benjamini-Hochberg correction (Storey and Tibshirani, 2003). The *qvalue* R package transforms the *p*-values into *q*-values and the user can control a specified value *α* of FDR by considering as candidates the SNPs with *q*-values smaller than *α*.

### Numerical computations

PCA is performed using a C routine that allows to compute scores and eigenvalues efficiently with minimum RAM access (Duforet-Frebourg et al., 2016). Computing the covariance matrix Ω is the most computationally demanding part. To provide a fast routine, we compute the *n* × *n* covariance matrix Ω instead of the much larger *p* × *p* covariance matrix. We compute the covariance Ω incrementally by adding small storable covariance blocks successively. Multiple linear regression is then solved directly by computing an explicit solution, written as a matrix product. Using the fact that the (*n*, *K*) score matrix *X* is orthogonal, the (*p*, *K*) matrix 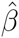 of regression coefficients is given by *G*^*T*^*X* and the (*n*, *p*) matrix of residuals is given by *G* − *XX^T^G*. The *z*-scores are then computed using the standard formula for multiple regression

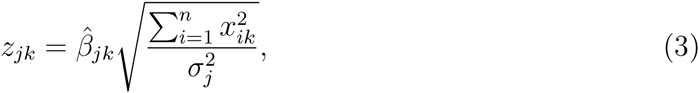

where 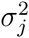 is an estimate of the residual variance for the *j*^th^ SNP, and *x_ik_* is the score of *k*^th^ principal component for the *i*^th^ individual.

### Missing data

Missing data should be accounted for when computing principal components and when computing the matrix of *z*-scores. There are many methods to account for missing data in PCA and we consider the pairwise covariance approach (Dray and Josse, 2015). It consists in estimating the covariance between each pair of individuals using only the markers that are available for both individuals. To compute *z*-scores, we account for missing data in formula (3). The term in the numerator 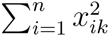 depends on the quantity of missing data. If there are no missing data, it is equal to 1 by definition of the scores obtained with PCA. As the quantity of missing data grows, this term and the *z*-score decrease such that it becomes more difficult to detect outlier markers.

### Pooled sequencing

When data are sequenced in pool, the Mahalanobis distance is based on the matrix of allele frequency computed in each pool instead of the matrix of *z*-scores.

## Materials and Methods

### Simulated data

We simulated SNPs under an island model, under a divergence model and we downloaded simulations of range expansion (Lotterhos and Whitlock, 2015). All data we simulated were composed of 3 populations, each of them containing 50 sampled diploid individuals (Table 2). SNPs were simulated assuming no linkage disequilibrium. SNPs with minor allele frequencies lower than 5% were discarded from the datasets. The mean *F_ST_* for each simulation was comprised between 5% and 10%. Using the simulations based on a island and a divergence model, we also created datasets composed of admixed individuals. We assumed that an instantaneous admixture event occurs at the present time so that all sampled individuals are the results of this admixture event. Admixed individuals were generated by drawing randomly admixture proportions using a Dirichlet distribution of parameter (*α*, *α*, *α*) (*α* ranging from 0.005 to 1 depending on the simulation).

**Table 2.** Summary of the simulations. The table above shows the average number of individuals, of SNPs, of adaptive markers and the total number of simulations per scenario.

|  | Individuals | SNPs | Adaptive SNPs | Simulations |
| --- | --- | --- | --- | --- |
| Island model | 150 | 472 | 27 | 35 |
| Divergence model | 150 | 3000 | 100 | 6 |
| Island model (hybrids) | 150 | 472 | 30 | 27 |
| Divergence model (hybrids) | 150 | 3000 | 100 | 9 |
| Range expansion | 1200 | 9999 | 99 | 6 |

### Island model

We used *ms* to create simulations under an island model (Fig SI1). We set a lower migration rate for the 50 adaptive SNPs compared to the 950 neutral ones to mimick diversifying selection (Bazin et al., 2010). For a given locus, migration from population *i* to *j* was specified by choosing a value of the effective migration rate that is set to *M*_neutral_ = 10 for neutral SNPs and to *M*_adaptive_ for adaptive ones. We simulated 35 datasets in the island model with different strengths of selection, where the strength of selection corresponds to the ratio *M*_neutral_/*M*_adaptive_ that varies from 10 to 1, 000. The *ms* command lines for neutral and adaptive SNPs are given by (*M*_adaptive_ = 0.01 and *M*_neutral_ = 10)

~~~
./ms 300 950 -s 1 -I 3 100 100 100 -ma x 10 10 10 x 10 10 10 x
./ms 300 50 -s 1 -I 3 100 100 100 -ma x 0.01 0.01 0.01 x 0.01 0.01 0.01 x
~~~

### Divergence model

To perform simulations under a divergence model, we used the package *simuPOP*, which is an individual-based population genetic simulation environment (Peng and Kimmel, 2005). We assumed that an ancestral panmictic population evolved during 20 generations before splitting into two subpopulations. The second subpopulation then split into subpopulations 2 and 3 at time *T* > 20. All 3 subpopulations continued to evolve until 200 generations have been reached, without migration between them (Figure SI1). A total of 50 diploid individuals were sampled in each population. Selection only occured in the branch associated with population 2 and selection was simulated by assuming an additive model (fitness is equal to 1 − 2*s*, 1 − *s*, 1 depending on the genotypes). We simulated a total of 3,000 SNPs comprising of 100 adaptive ones for which the selection coefficient is of *s* = 0.1.

### Range expansion

We downloaded in the *Dryad Digital Repository* six simulations of range expansion with two glacial refugia (Lotterhos and Whitlock, 2015). Adaptation occurred during the recolonization phase of the species range from the two refugia. We considered six different simulated data with 30 populations and a number of sampled individual per location that varies from 20 to 60.

### Parameter settings for the different software

When using *hapflk*, we set *K* =1 that corresponds to the computation of the *FLK* statistic. When using *BayeScan* and *OutFLANK*, we used the default parameter values. For *sNMF*, we used *K* = 3 for the island and divergence model and *K* = 5 for range expansion as indicated by the cross-entropy criterion. The regularization parameter of *sNMF* was set to *α* = 1000. For *sNMF* and *hapflk*, we used the genomic inflation factor to recalibrate *p*-values. When using population-based methods with admixed individuals, we assigned each individual to the population with maximum amount of ancestry.

## Results

### Choosing the number of principal components

We evaluate Cattell’s graphical rule to choose the number of principal components. For the island and divergence model, the choice of *K* is evident (Figure 1). For *K* ≥ 3, the eigenvalues follow a straight line. As a consequence, Cattell’s rule indicates *K* = 2, which is expected because there are 3 populations (Patterson et al., 2006). For the model of range expansion, applying Cattell’s rule to choose *K* is more difficult (Figure 1). Ideally, the eigenvalues that correspond to random variation lie on a straight line whereas the ones corresponding to population structure depart from the line. However, there is no obvious point at which eigenvalues depart from the straight line. Choosing a value of *K* between 5 and 8 is compatible with Cattell’s rule. Using the package *qvalue* to control 10% of FDR, we find that the actual proportion of false discoveries as well as statistical power is weakly impacted when varying the number of principal components from *K* = 5 to *K* = 8 (Figure SI2).

**Figure 1.**
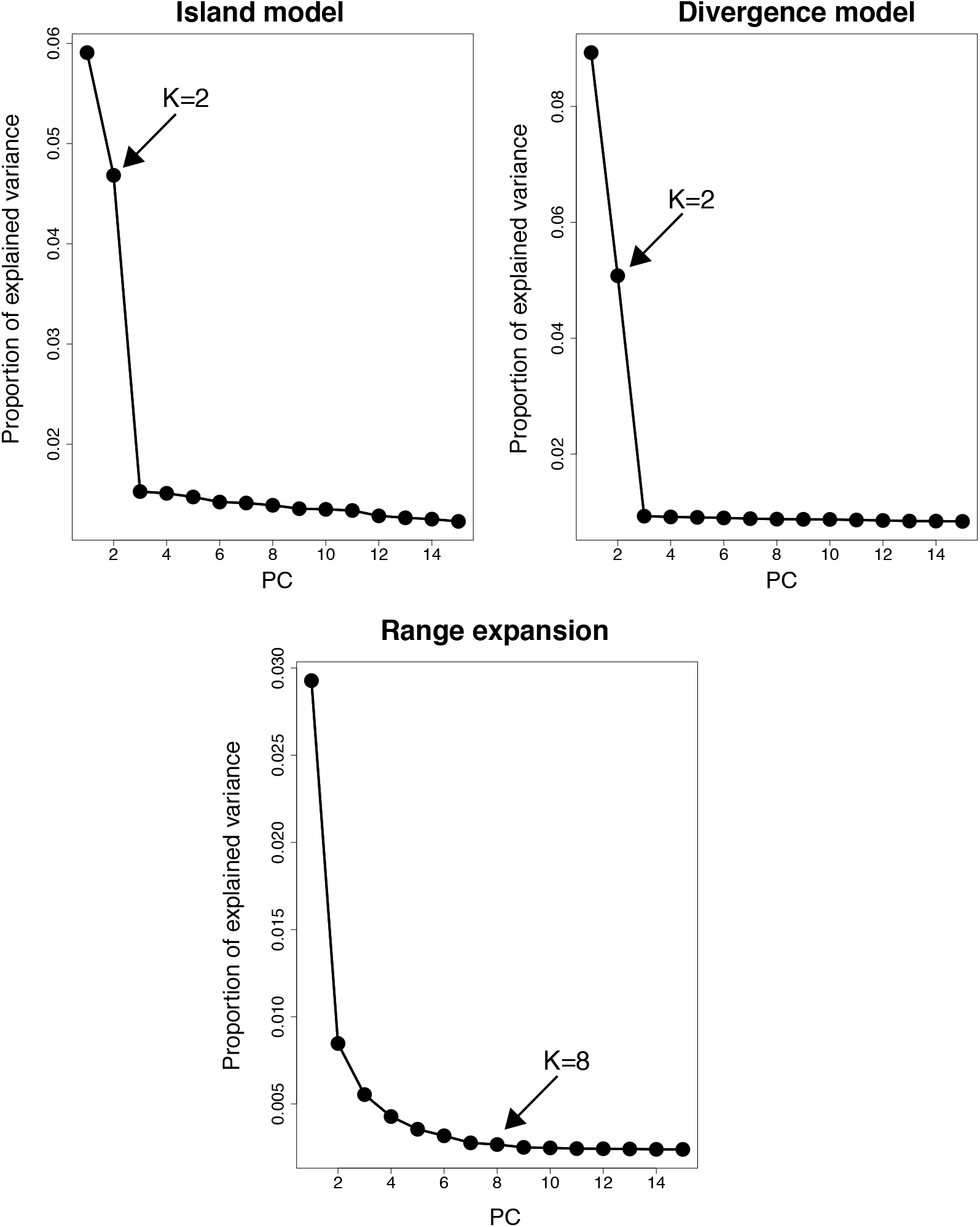
Determining *K* with the screeplot. To choose *K*, we recommend to use Cattell’s rule that states that components corresponding to eigenvalues to the left of the straight line should be kept. According to Cattell’s rule, the eigenvalues that correspond to random variation lie on the straight line whereas the ones corresponding to population structure depart from the line. For the island and divergence model, the choice of *K* is evident. For the model or range expansion, a value of *K* between 5 and 8 is compatible with Cattell’s rule.

### An example of genome scans performed with *pcadapt*

To provide an example of results, we apply *pcadapt* with *K* = 6 in the model of range expansion. Population structure captured by the first 2 principal components is displayed in Figure 2. *P*-values are well calibrated because they are distributed as a mixture of a uniform distribution and of a peaky distribution around 0, which corresponds to outlier loci (Figure 2). Using a FDR threshold of 10% with the *qvalue* package, we find 122 outliers among 10, 000 SNPs, resulting in 23% actual false discoveries and a power of 95%.

**Figure 2.**
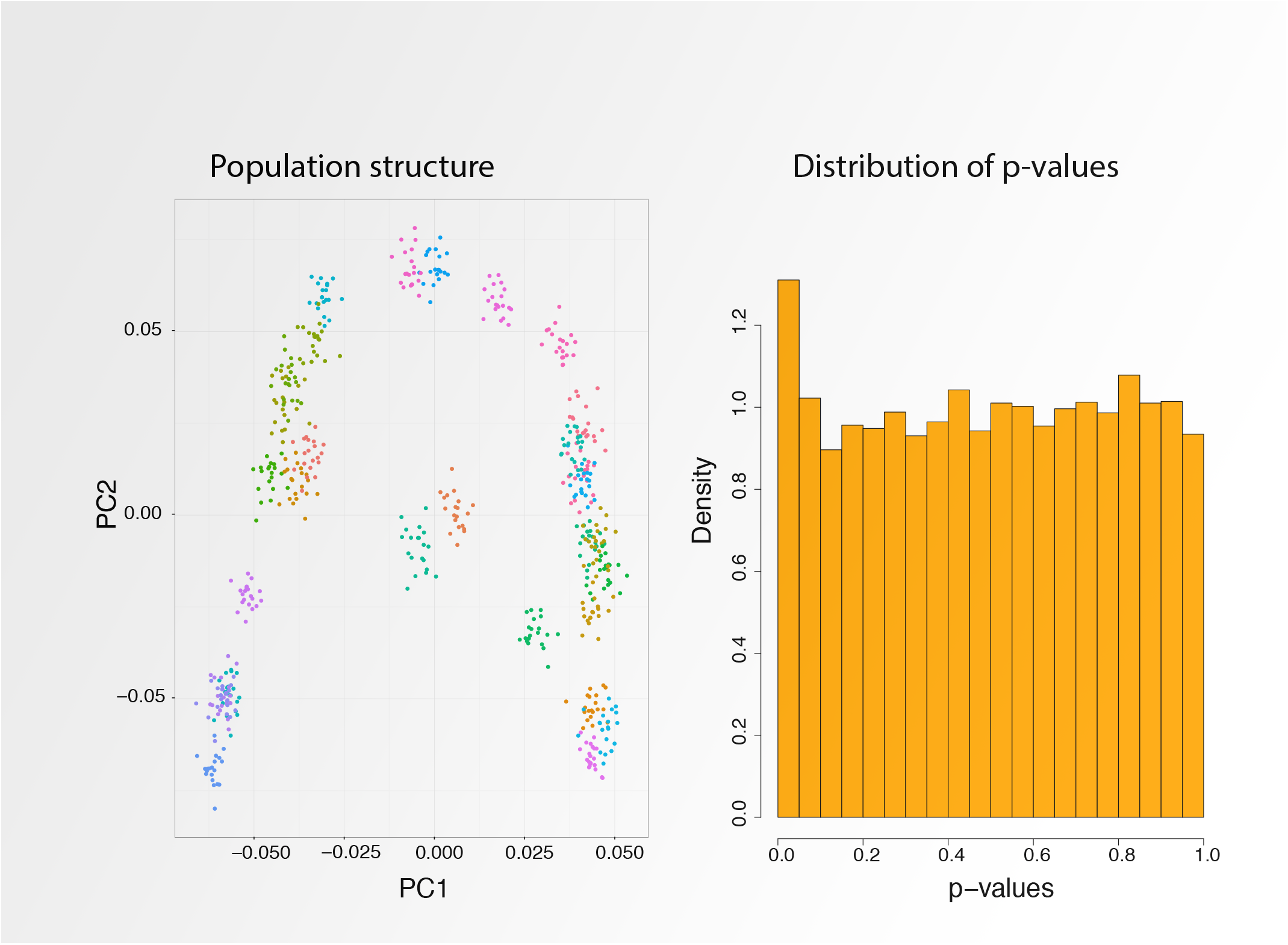
Population structure (first 2 principal components) and distribution of p-value obtained with *pcadapt* for a simulation of range expansion. *P*-values are well calibrated because they are distributed as a mixture of a uniform distribution and of a peaky distribution around 0, which corresponds to outlier loci. In the left panel, each color corresponds to individuals sampled from the same population.

### Control of the false discovery rate

We evaluate to what extent using the packages *pcadapt* and *qvalue* control a FDR set at 10% (Figure 3). All SNPs with a *q*-value smaller than 10% were considered as candidate SNPs. For the island model, we find that the proportion of false discoveries is 8% and it increases to 10% when including admixture. For the divergence model, the proportion of false discoveries is 11% and it increases to 22% when including admixture. The largest proportion of false discoveries is obtained under range expansion and is equal to 25%.

**Figure 3.**
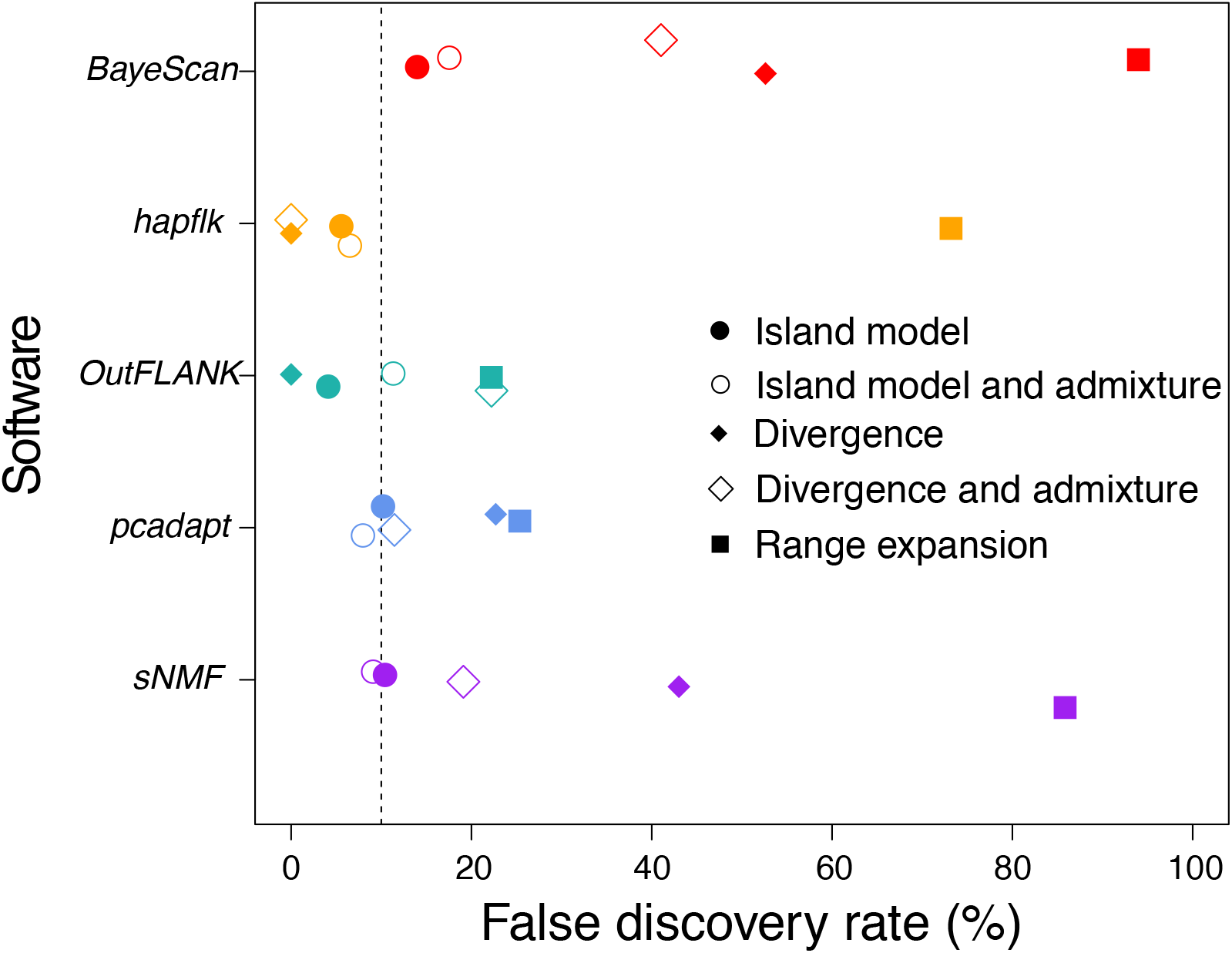
Control of the FDR for different software of genome scans. We find that the median proportion of false discoveries is around the nominal FDR set at 10% (6% for *hapflk*, 11% for both *OutFLANK* and *pcadapt*, and 19% for *sNMF*) with the exception of *BayeScan* that generates 41% of false discoveries.

We then evaluate the proportion of false discoveries obtained with *BayeScan*, *hapflk*, *OutFLANK*, and *sNMF* (Figure 3). We find that *hapflk* is the most conservative approach (FDR = 6%) followed by *OutFLANK* and *pcadapt* (FDR = 11%). The software *sNMF* is more liberal (FDR = 19%) and *BayeScan* generates the largest proportion of false discoveries (FDR = 41%). When not recalibrating the *p*-values of *hapflk*, we find that the test is even more conservative (results not shown). For all software, the range expansion scenario is the one that generates the largest proportion of false discoveries. Proportion of false discoveries under range expansion ranges from 22% (*OutFLANK*) to 93% (*BayeScan*).

### Statistical power

To provide a fair comparison between methods and software, we compare statistical power for equal values of the observed proportion of false discoveries. Then we compute statistical power averaged over observed proportion of false discoveries ranging from 0% to 50%.

We first compare statistical power obtained with the different statistical methods that have been implemented in *pcadapt* (Table 1). For the island model, Bayes factor, communality statistic and Mahalanobis distance have similar power (Figure 4). For the divergence model, the power obtained with Mahalanobis distance is 20% whereas the power obtained with the communality statistic and with the Bayes factor is respectively 4% and 2% (Figure 4). Similarly, for range expansion, the power obtained with Mahalanobis distance is 46% whereas the power obtained with the communality statistic and with the Bayes factor is 34% and 13%. We additionally investigate to what extent increasing sample size in each population from 20 to 60 individuals affects power. For range expansion, the power obtained with the Mahalanobis distance hardly changes ranging from 44% to 47%. However, the power obtained with the other two statistics changes importantly. The power obtained with the communality statistic increases from 27% to 39% when increasing the sample size and the power obtained with the Bayes factor increases from 0% to 44%.

**Figure 4.**
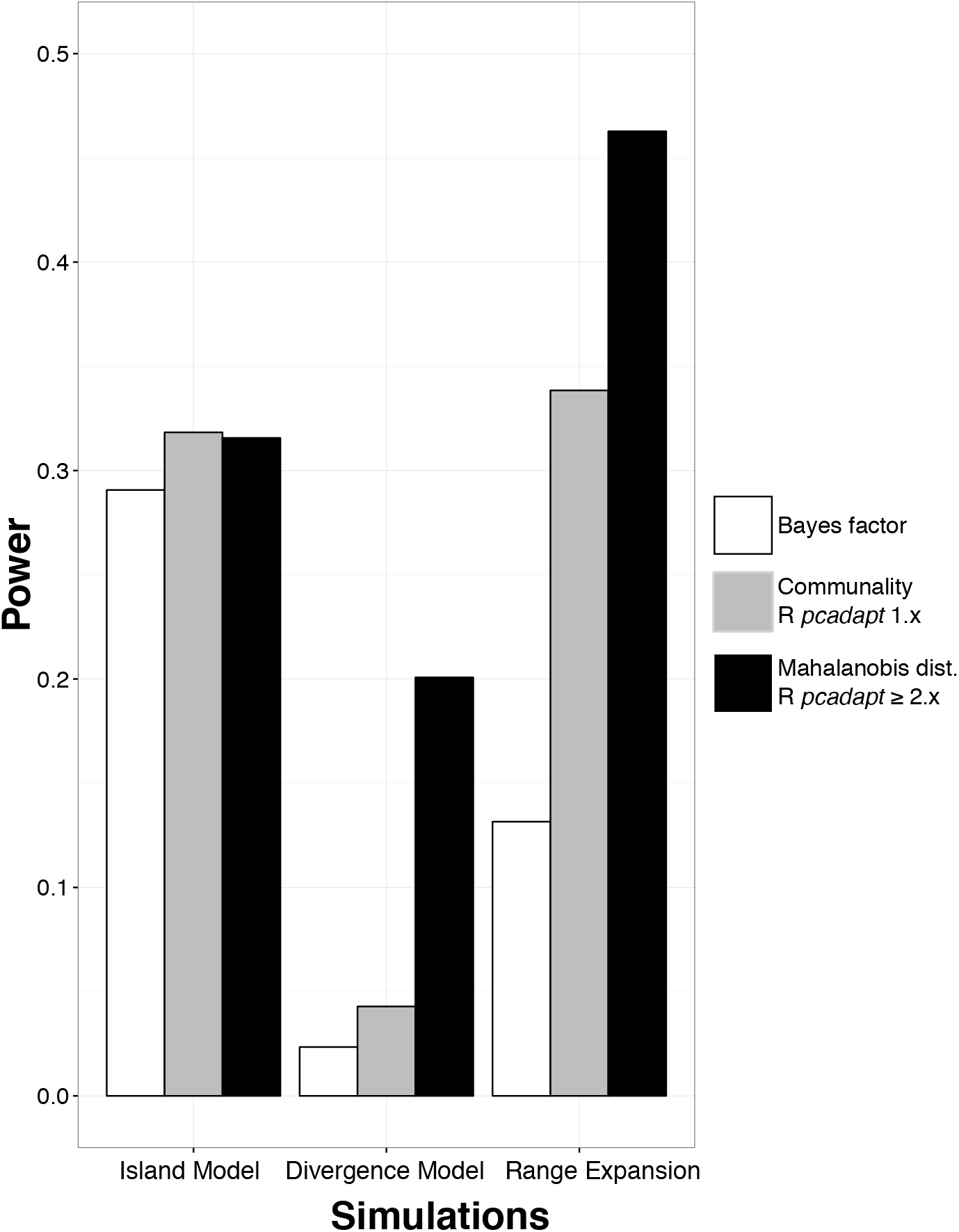
Comparison of statistical power for the different test statistics that have been implemented in *pcadapt* (Table 1). Bayes factors corresponds to the test statistics implemented in the Bayesian version of *pcadapt* (Duforet-Frebourg et al., 2014); the communality statistic was the default statistic in version 1.x of the R package *pcadapt* (Duforet-Frebourg et al., 2016), and Mahalanobis distances are available since the release of the 2.0 version of the package. When there is hierarchical population structure (divergence model and range expansion), the Mahalanobis distance provides more powerful genome scans compared to the test statistic previously implemented in *pcadapt*. The abbreviation dist. stands for distance. Statistical power is averaged over the observed proportion of false discoveries (ranging between 0% and 50%).

Then we describe our comparison of software for genome scans. For the simulations obtained with the island model where there is no hierarchical population structure, the statistical power is similar for all software (Figure SI3 and SI4). Including admixed individuals hardly changes their statistical power (Figure SI3).

Then, we compare statistical power in a divergence model where adaptation took place in one of the external branches of the population divergence tree. The software *pcadapt* and *hapflk*, which account for hierarchical population structure, as well as *BayeScan* are the most powerful in that setting (Figure 5 and Figure SI5). The values of power in decreasing order are of 23% for *BayeScan*, of 20% for *pcadapt*, of 17% for *hapflk*, of 7% for *sNMF* and of 1% for *OutFLANK*. When including admixed individuals, the power of *hapflk* and of *pcadapt* hardly decreases whereas the power of *BayeScan* decreases to 6% (Figure 5).

**Figure 5.**
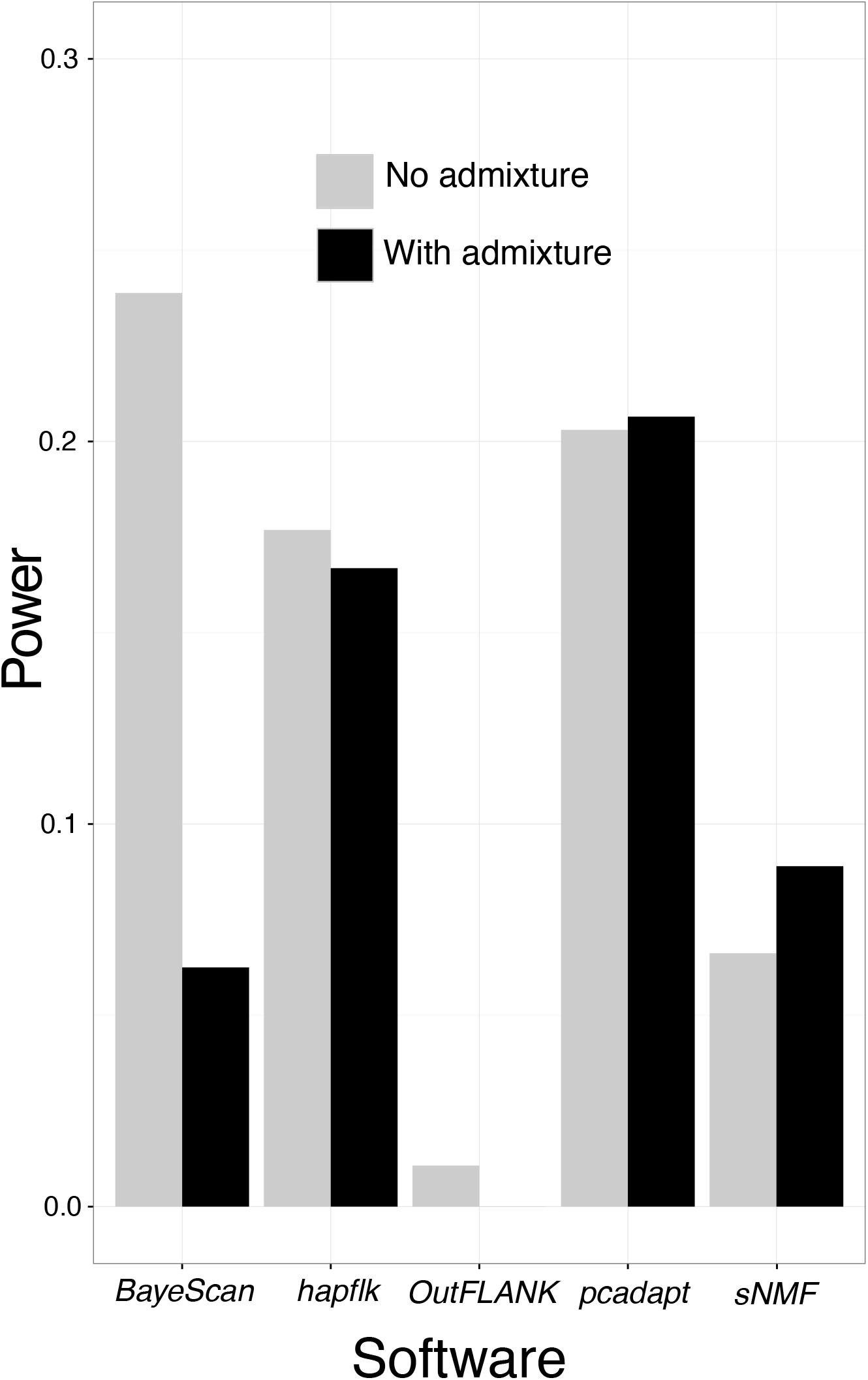
Statistical power averaged over the expected proportion of false discoveries (ranging between 0% and 50%) for the divergence model with 3 populations. We assume that adaptation took place in an external branch that follows the most recent population divergence event.

The last model we consider is the model of range expansion. The package *pcadapt* is the most powerful approach in this setting (Figure 6 and SI6). Other software also discover many true positive loci with the exception of *BayeScan* that provides no true discovery when the observed FDR is smaller than 50% (Figure 6 and SI6). The values of power in decreasing order are of 46% for *pcadapt*, of 41% for *hapflk*, of 37% for *OutFLANK*, of 30% for *sNMF* and of 0% for *BayeScan*.

**Figure 6.**
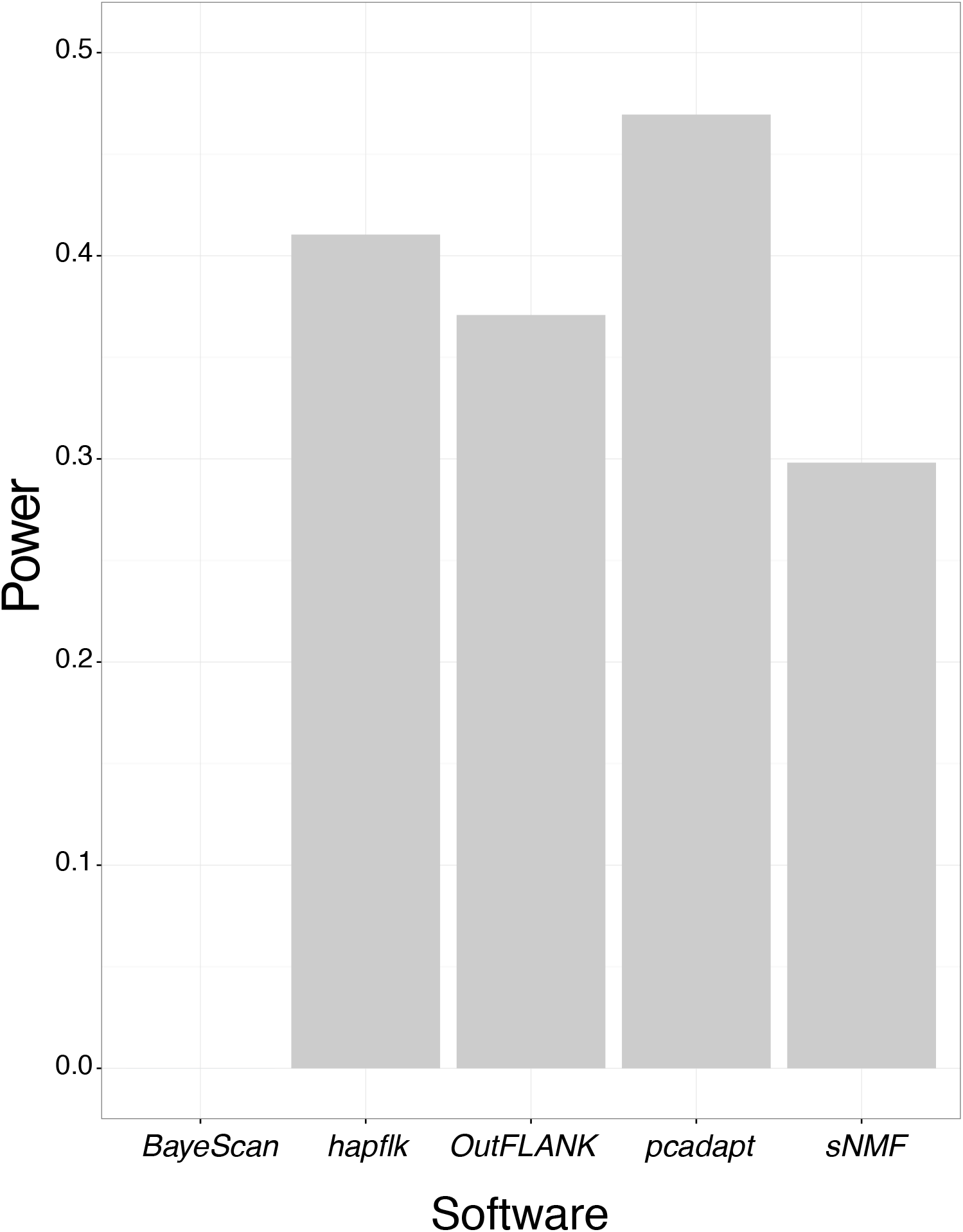
Statistical power averaged over the expected proportion of false discoveries (ranging between 0% and 50%) for a range expansion model with two refugia. Adaptation took place during the recolonization event.

### Running time of the different software

Last, we compare the software running times. The characteristics of the computer we used to perform comparisons are the following: OSX El Capitan 10.11.3, 2,5 GHz Intel Core i5, 8 Go 1600 MHz DDR3. We discard *BayeScan* as it is too time consuming compared to other software. For instance, running *BayeScan* on a genotype matrix containing 150 individuals and 3, 000 SNPs takes 9 hours whereas it takes less than one second with *pcadapt*. The different software were run on genotype matrices containing 300 individuals and from 500 to 50, 000 SNPs. *OutFLANK* is the software for which the runtime increases the most rapidly with the number of markers. *OutFLANK* takes around 25 minutes to analyse 50,000 SNPs (Figure SI7). For the other 3 software (*hapflk*, *pcadapt*, *sNMF*), analyzing 50, 000 SNPs takes less than 3 minutes.

## Discussion

The R package *pcadapt* implements a fast method to perform genome scans with next generation sequencing data. It can handle datasets where population structure is continuous or datasets containing admixed individuals. It can handle missing data as well as pooled sequencing data. The 2.0 and later versions of the R package implements a robust Mahalanobis distance as a test statistic. When hierarchical population structure occurs, Mahalanobis distance provides more powerful genome scans compared to the communality statistic that was implemented in the first version of the package (Duforet-Frebourg et al., 2016). In the divergence model, adaptation occurs along an external branch of the divergence tree that corresponds to the second principal component. When outlier SNPs are not related to the first principal component, the Mahalanobis distance provides a better ranking of the SNPs compared to the communality statistic.

Simulations show that the R package *pcadapt* compares favorably to other software of genome scans. When data were simulated under an island model, population structure is not hierarchical because genetic differentiation is the same for all pairs of populations. Statistical power and control of the FDR were similar for all software. In presence of hierarchical population structure (divergence model) where genetic differentiation varies between pairs of populations, the ranking of the SNPs is software dependent. The software *pcadapt* and *hapflk* provide the most powerful scans whether or not simulations include admixed individuals. *OutFLANK* implements a *F_ST_* statistic and because adaptation does not correspond to the most differentiated populations, it fails to capture adaptive SNPs (Figure 5) (Bonhomme et al., 2010; Duforet-Frebourg et al., 2016). *BayeScan* does not assume equal differentiation between all pairs of populations, which may explain why it has a good statistical power for the divergence model. However its statistical power is severely impacted by the presence of admixed individuals because its power decreases from 24% to 6% (Figure 5). Understanding why *BayeScan* is severely impacted by admixture is out of the scope of this paper. In the range expansion model, *BayeScan* returns many null *q*-values (between 376 and 809 SNPs out of 9, 899 neutral and 100 adaptive SNPs) such that the observed FDR is always larger than 50%. Overall, we find that *pcadapt* and *hapflk* provides comparable statistical power. Compared to other software, they provide optimal or near optimal ranking of the SNPs in different scenarios including hierarchical population structure and admixed individuals. The main difference between the two software concerns the control of the FDR because *hapflk* is found to be more conservative.

Because NGS data become more and more massive, careful numerical implementation is crucial. There are different options to implement PCA and *pcadapt* uses a numerical routine based on the computation of the covariance matrix Ω. The algorithmic complexity to compute the covariance matrix is proportional to *pn*^2^ where p is the number of markers and *n* is the number of individuals. The computation of the first *K* eigenvectors of the covariance matrix Ω has a complexity proportional to *n*^3^. This second step is usually more rapid than the computation of the covariance because the number of markers is usually large compared to the number of individuals. In brief, computing the covariance matrix Ω is by far the most costly operation when computing principal components. Although we have implemented PCA in *C* to obtain fast computations, an improvement in speed could be envisioned for future versions. When the number of individuals becomes large (e.g. *n* ≥ 10, 000), there are faster algorithms to compute principal components (Halko et al., 2011; Abraham and Inouye, 2014). In addition to running time, numerical implementations also impact the effect of missing data on principal components (Dray and Josse, 2015). Achieving a good tradeoff between fast computations and accurate evaluation of population structure in the face of large amount of missing data is a challenge for modern numerical methods in molecular ecology.

## Acknowledgements

This work has been supported by the LabEx PERSYVAL-Lab (ANR-11-LABX-0025-01) and the ANR AGRHUM project (ANR-14-CE02-0003-01).

## Data Accessibility

Island and divergence model data: doi:10.5061/dryad.8290n

Range expansion simulated data: doi:10.5061/dryad.mh67v. Files:

~~~
2R_R30_1351142954_453_2_NumPops=30_Numlnd=20
2R_R30_1351142954_453_2_NumPops=30_Numlnd=60
2R_R30_1351142970_988_6_NumPops=30_Numlnd=20
2R_R30_1351142970_988_6_NumPops=30_Numlnd=60
2R_R30_1351142986_950_10_NumPops=30_NumInd=20
2R_R30_1351142986_950_10_NumPops=30_NumInd=60
~~~

---

**Figure SI1.**
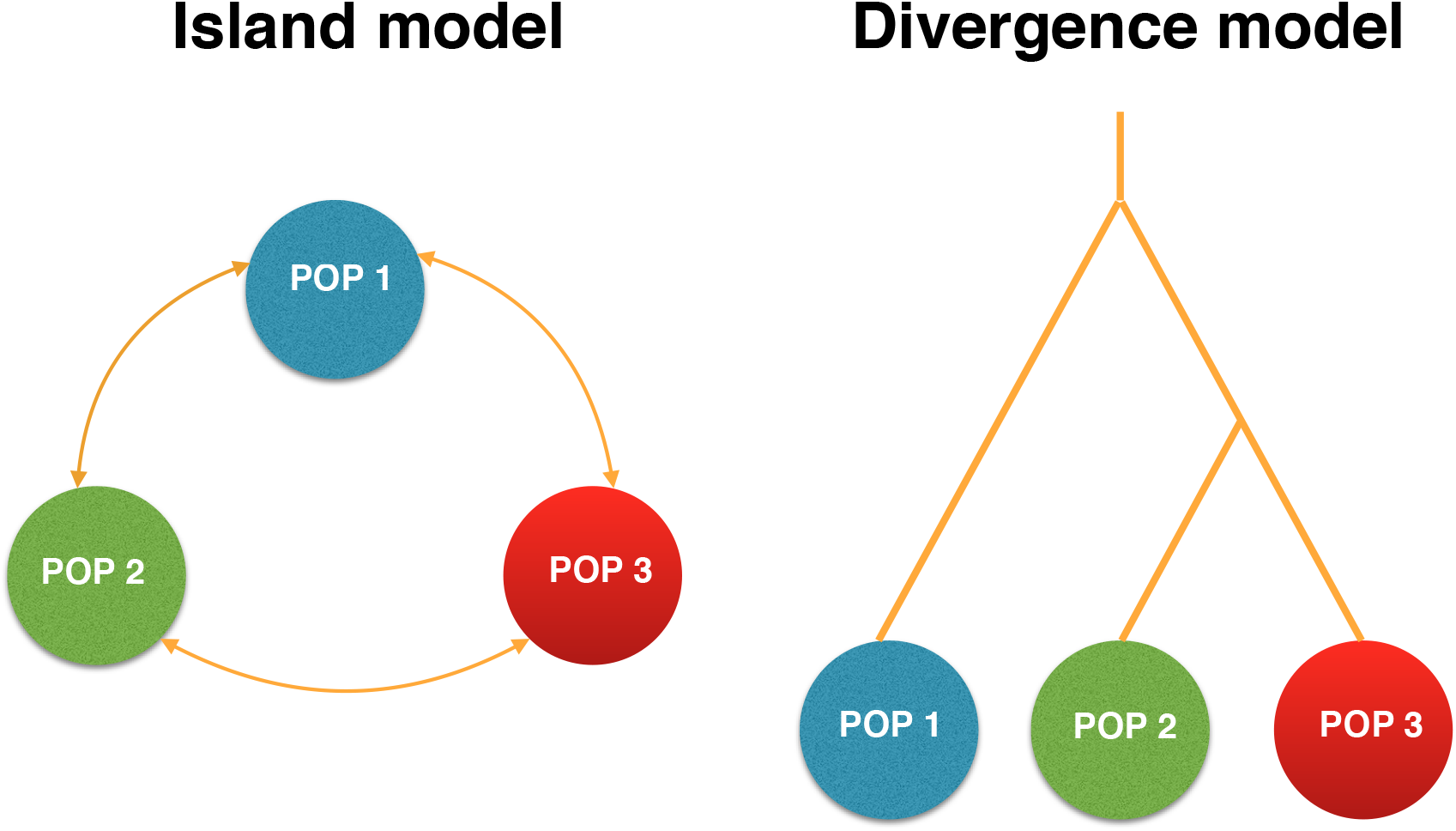
Schematic description of the island and divergence model. For the island model, adaptation occurs simultaneously in each population. For the island model, adaptation takes place in the branch leading to the second population.

**Figure SI2.**
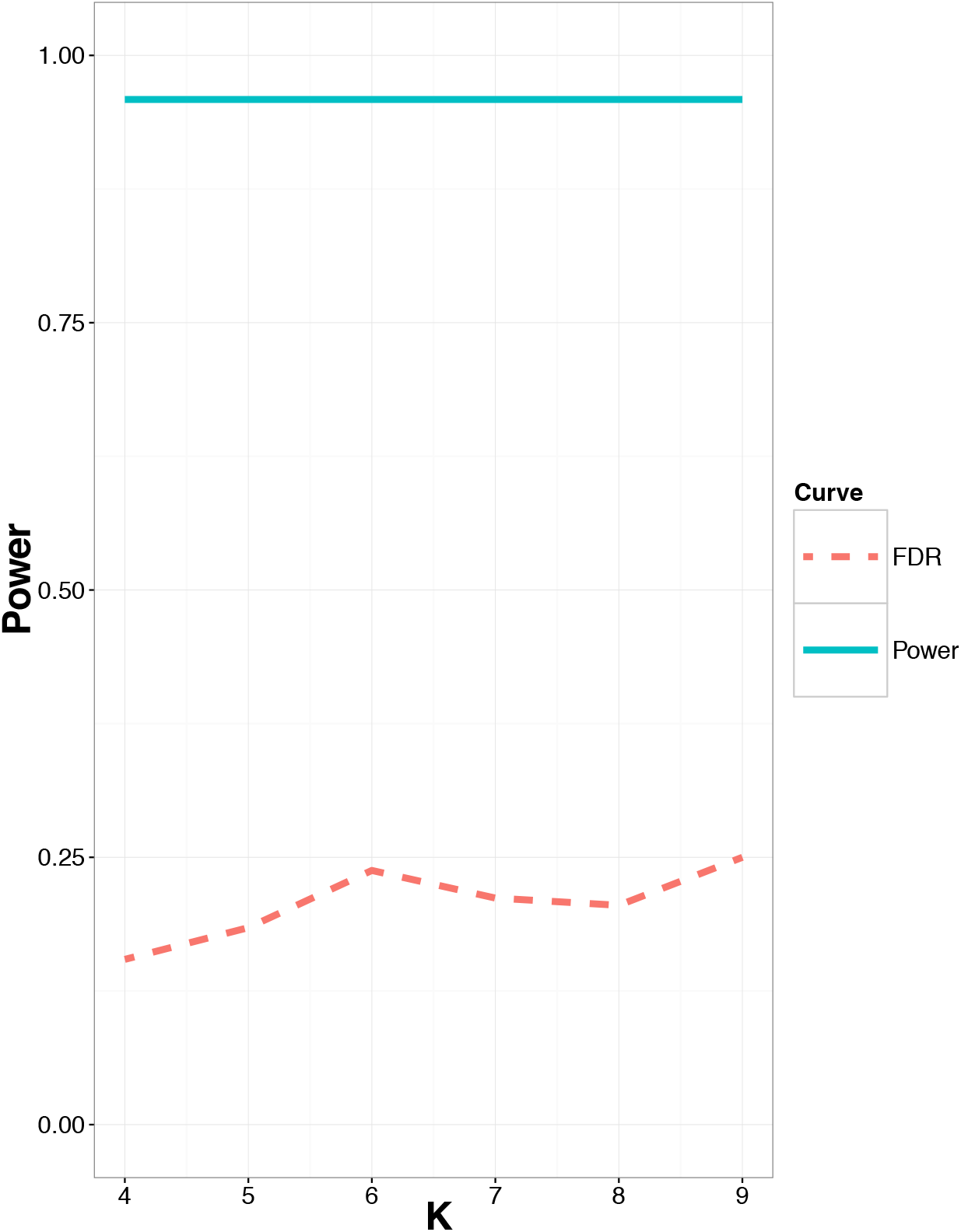
Proportion of false discoveries and statistical power as a function of the number of principal components in a model of range expansion.

**Figure SI3.**
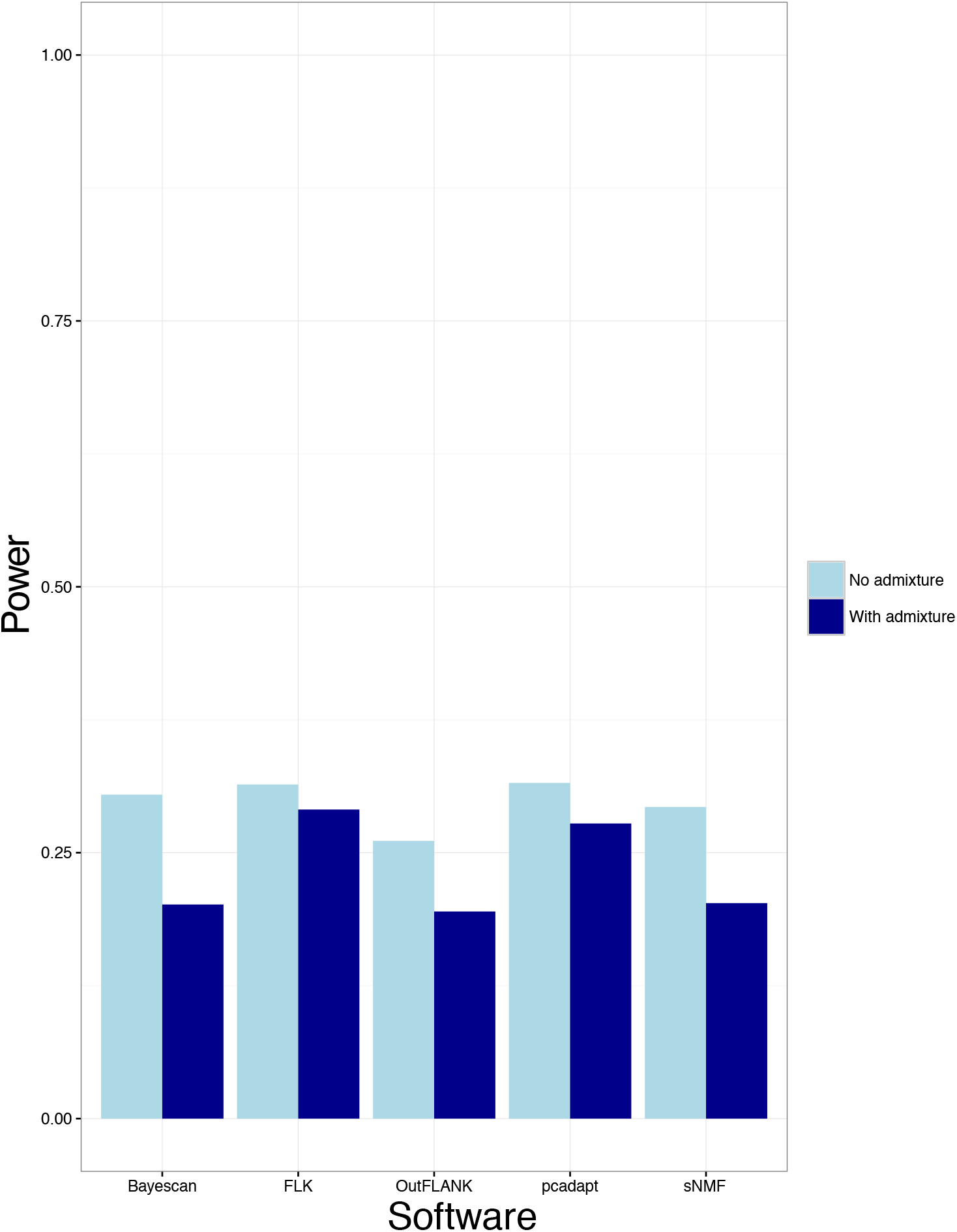
Statistical power averaged over the expected proportion of false discoveries (ranging between 0% and 50%) for the island model.

**Figure SI4.**
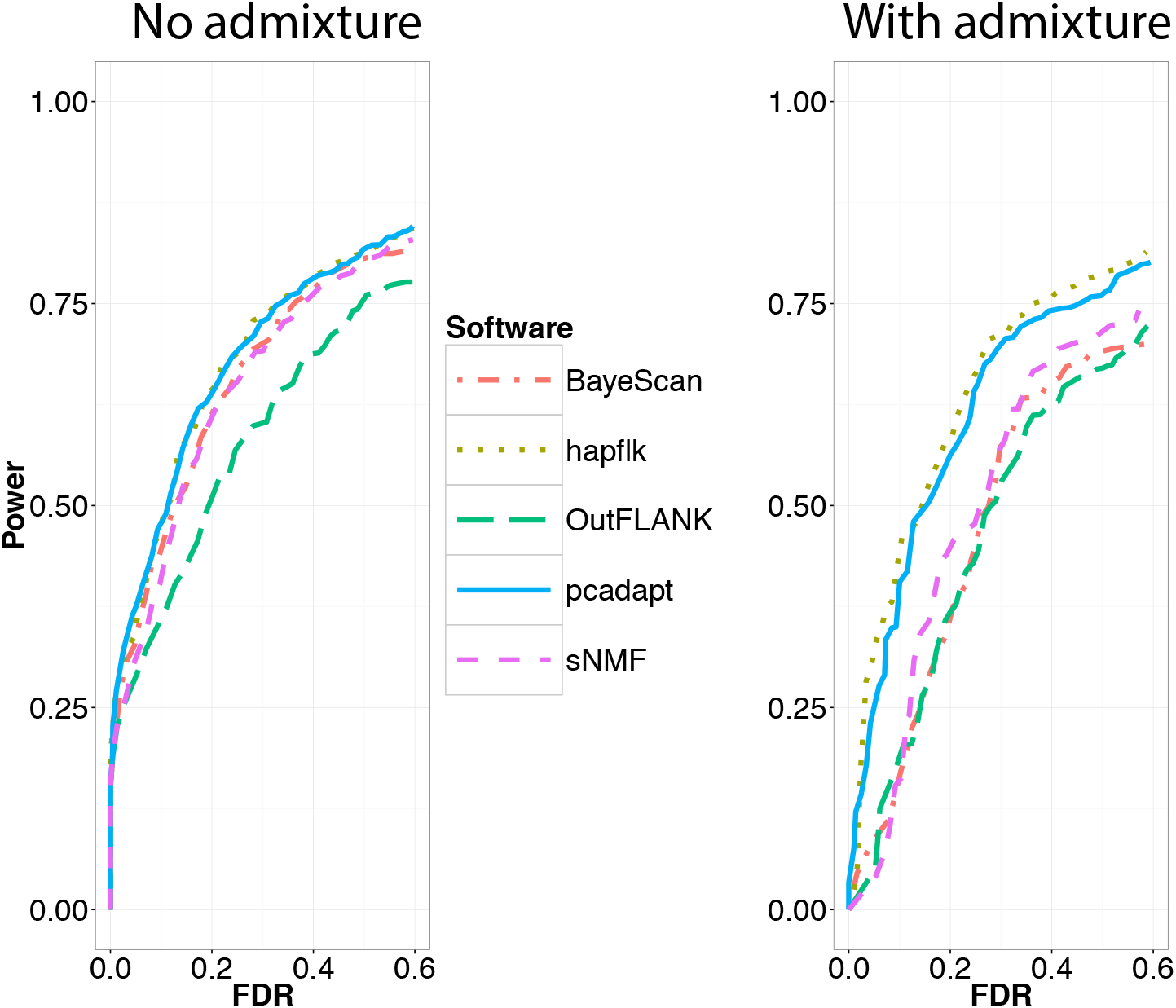
Statistical power as a function of the proportion of false discoveries for the island model.

**Figure SI5.**
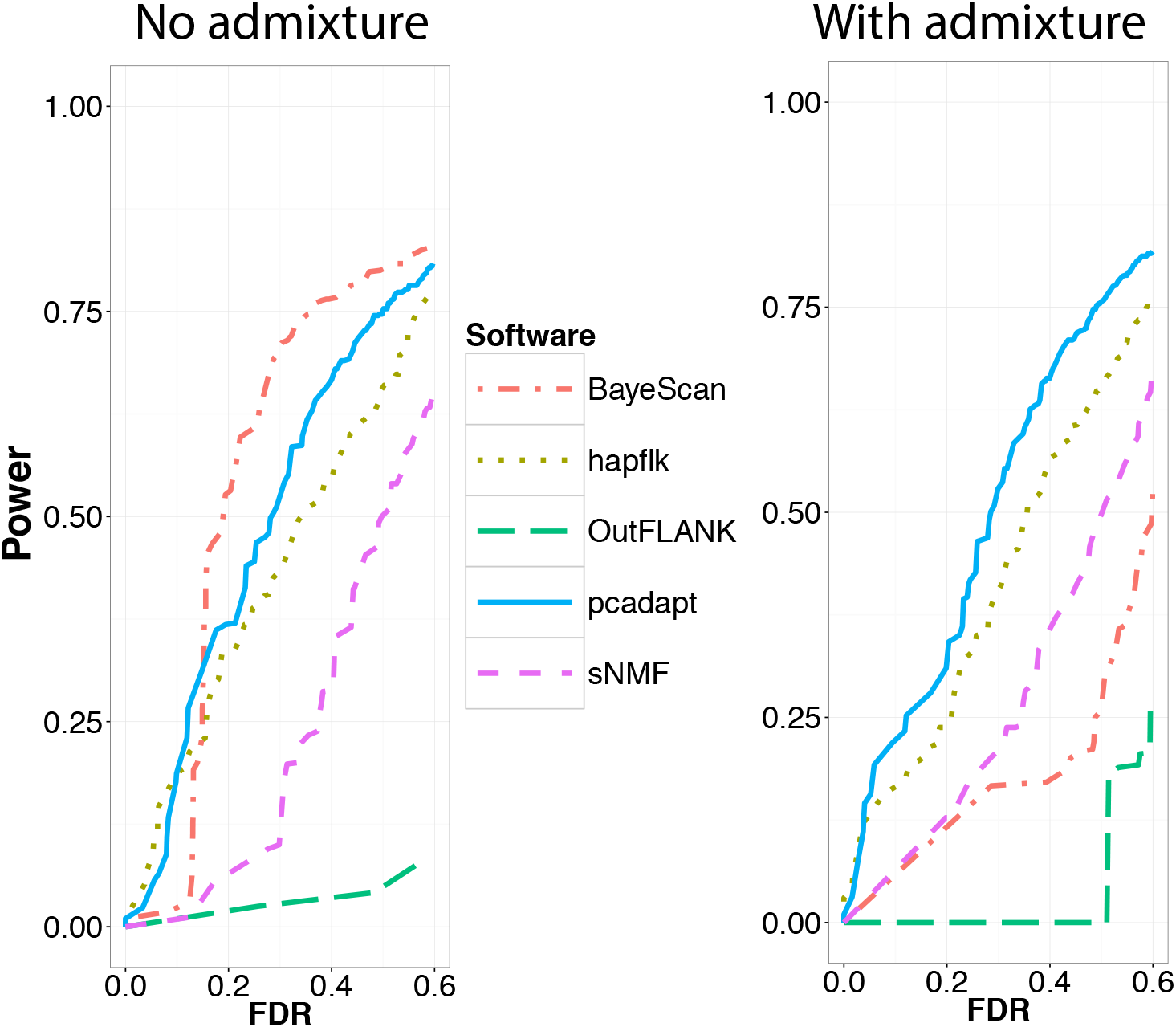
Statistical power as a function of the proportion of false discoveries for the divergence model.

**Figure SI6.**
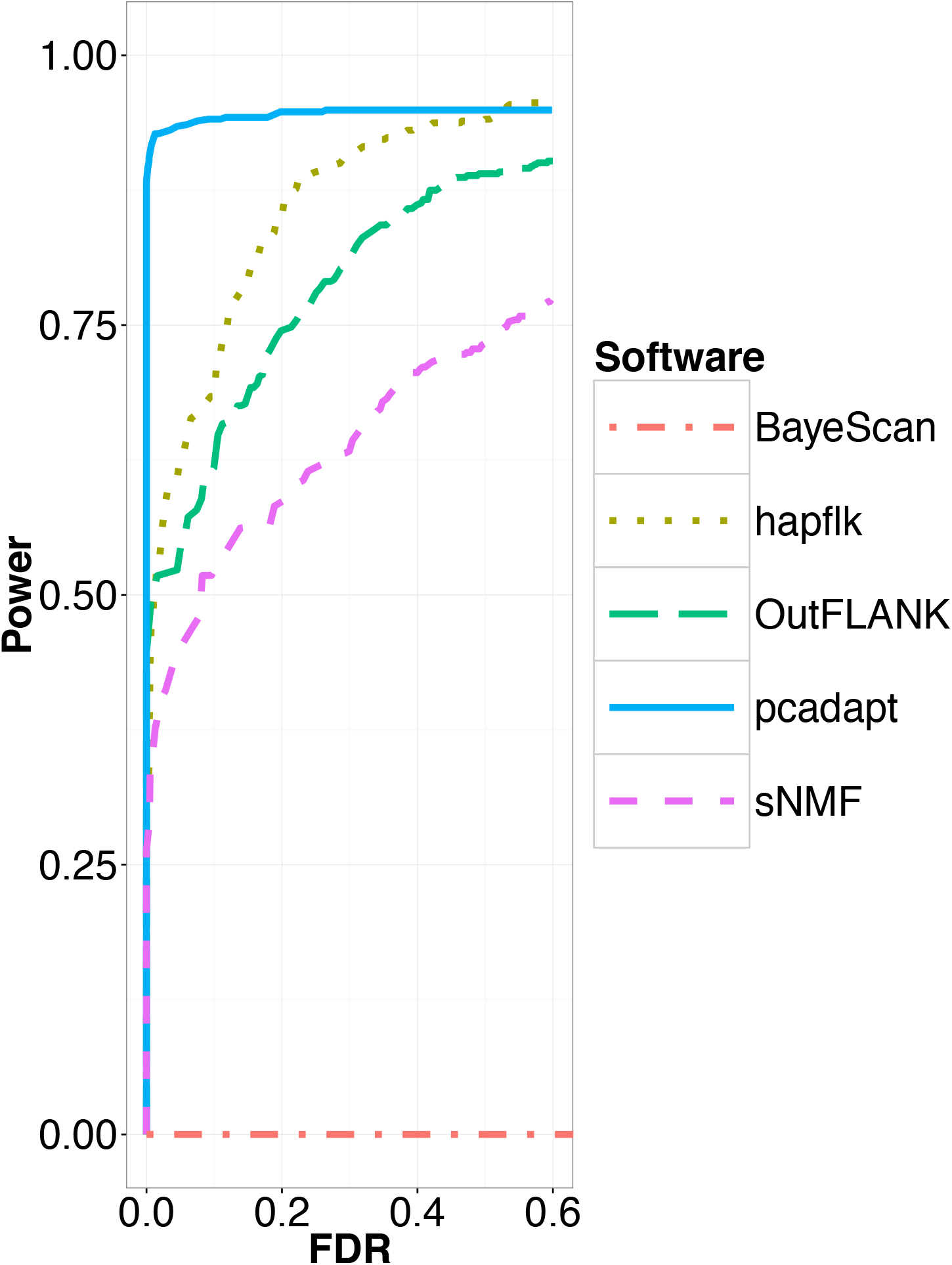
Statistical power as a function of the proportion of false discoveries for the model of range expansion.

**Figure SI7.**
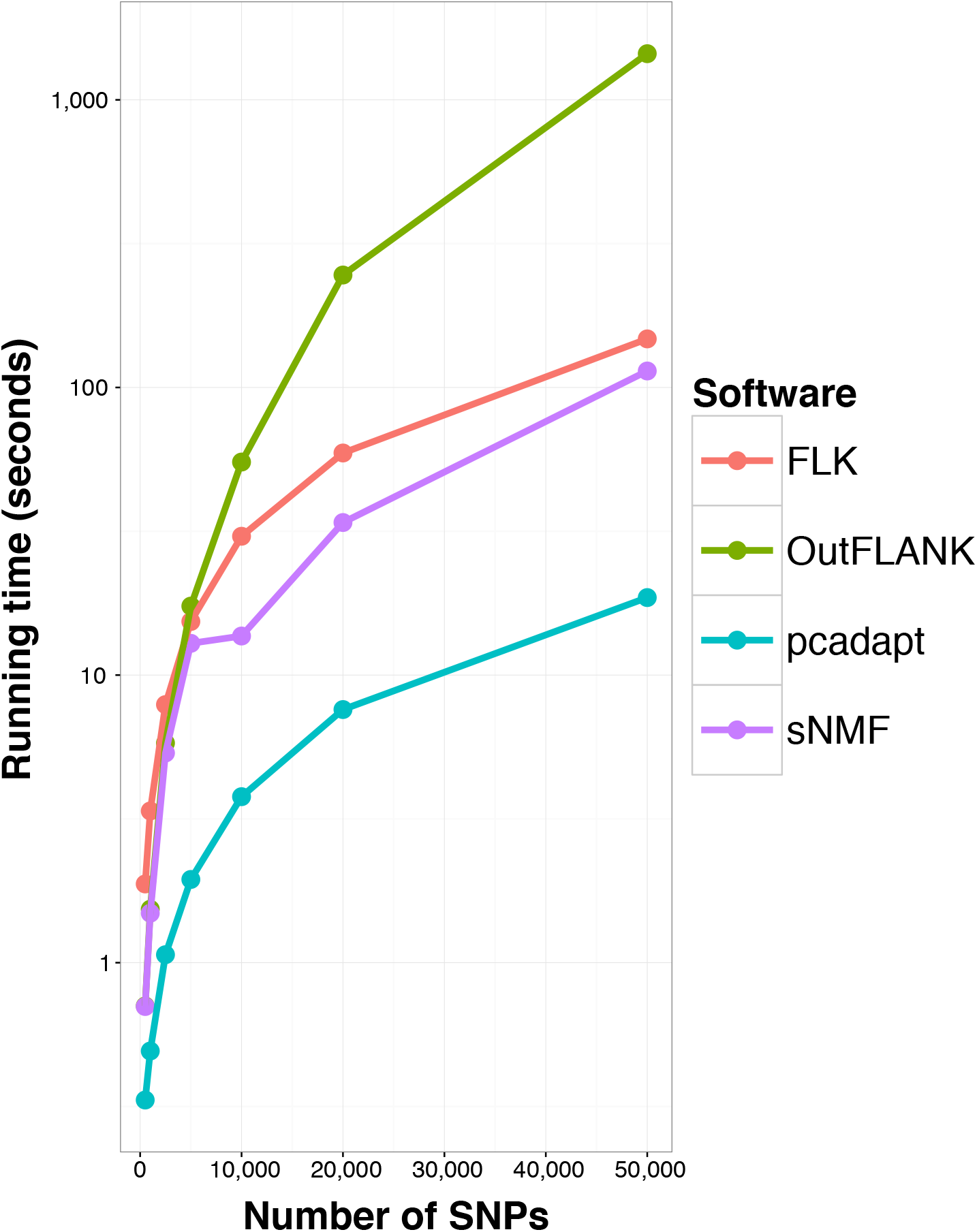
Running times of the different software. The different software were run on genotype matrices containing 300 individuals and from 500 to 50,000 SNPs. The characteristics of the computer we used to perform comparisons is the following: OSX El Capitan 10.11.3, 2,5 GHz Intel Core i5, 8 Go 1600 MHz DDR3.

